# Engineered caspases directly rewire mutant Ras to cell death

**DOI:** 10.64898/2026.08.06.743376

**Authors:** Lukas Moeller, Andrew C. Lu, Kevin Ho, Evan Zhang, Michael B. Elowitz

## Abstract

As central executioners of cell death, caspases that activate exclusively in diseased cells would provide powerful and specific therapeutic agents. Natural caspase regulation exhibits two universal features that facilitate the engineering of such caspases: proximity-induced subunit assembly and modular separation of substrate recruitment from catalysis. Here, we take advantage of these features to engineer “Raspases,” split effector caspases that conditionally reconstitute active complexes upon detection of mutant Ras, an oncogene altered in roughly a quarter of all cancers. When delivered as mRNA in lipid nanoparticles, Raspases selectively eliminate Ras-mutant human cancer lines while sparing wild-type cells. The system is built entirely from human protein domains, can be encoded as a single polyprotein, and can be adapted to trigger pyroptosis. Critically, Raspases match or exceed the potency of alternative Ras-targeting interventions in vitro. These results establish retargeted caspases as a generalizable sense-and-kill platform for selective elimination of diseased cells.

## Introduction

Across the animal kingdom, caspases are the key regulators and executioners of programmed cell death ^1^. As proteases, caspases break down cells from within, cleaving cytoskeletal and nuclear proteins and inducing genome fragmentation ^2^. This regulated form of cell death sculpts embryonic development, eliminates the surplus neurons generated during nervous-system maturation, and clears the billions of damaged cells in our bodies every day^1–4^. The human genome encodes roughly a dozen caspases in two classes^5,6^. Apoptotic caspases include initiators (caspase-2, -8, -9, -10), which receive and process death signals, and executioners (caspase-3, -6, -7), which carry them out ^5,6^. Inflammatory caspases (caspase-1, -4, -5) drive cytokine maturation and execute pyroptosis, the lytic form of programmed cell death ^7^.

Despite their power, efforts to harness caspases therapeutically have remained limited. Current approaches, such as small-molecule caspase inhibitors, peptide mimetics, and pan-caspase activators, share two fundamental limitations ^8–11^. First, they lack conditionality, activating or inhibiting caspase function in both healthy and diseased cells^8–11^. Second, by acting on conserved active-site residues, they often hit multiple paralogs, risking simultaneous perturbation of inflammatory, apoptotic, and homeostatic caspase programs^8,12^. The ability to programmably retarget specific caspases toward disease-specific signals could overcome both limitations and unlock the full therapeutic potential of caspases (**Figure 1A**).

**Figure 1.**
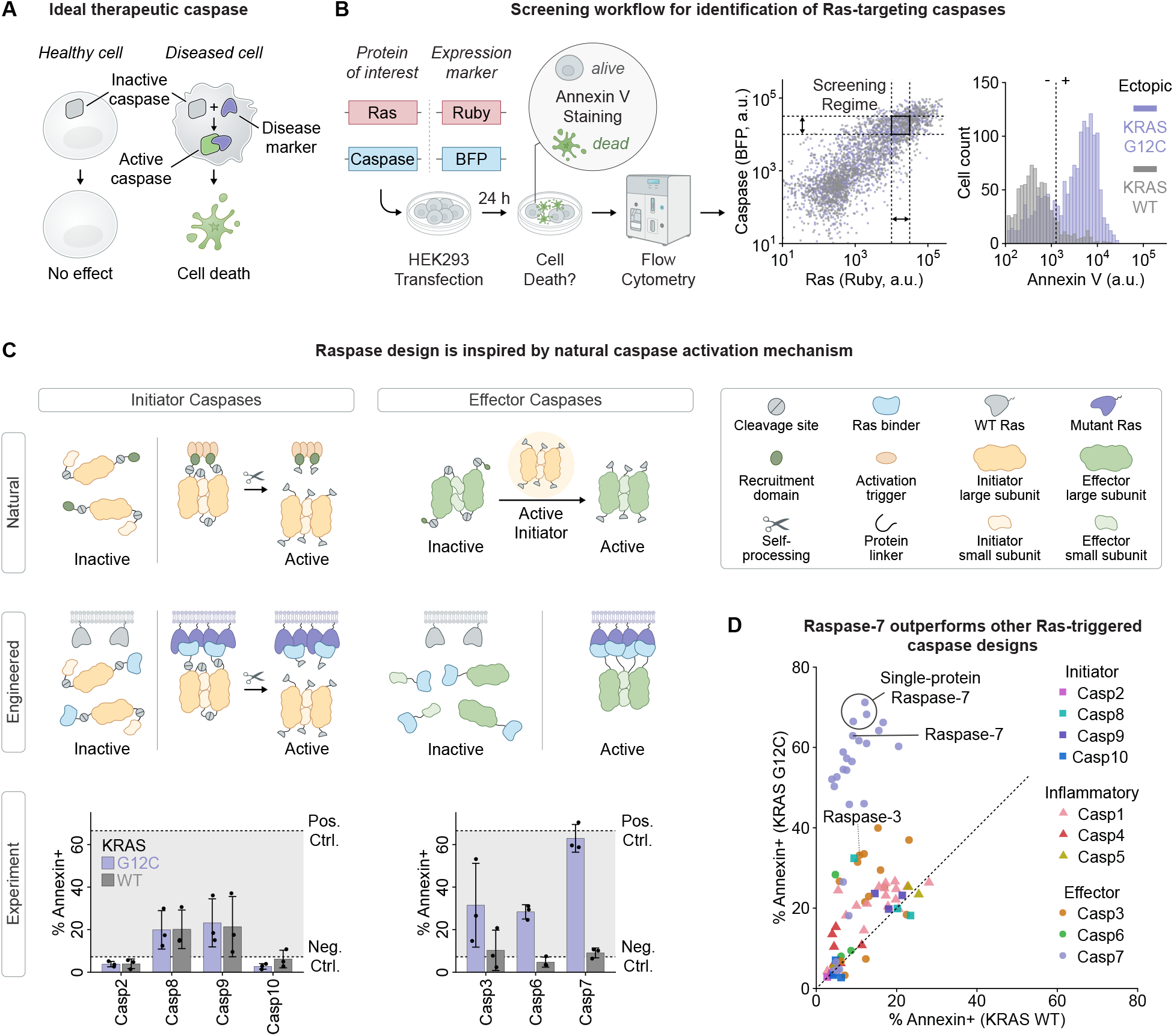
Mutant Ras-dependent reconstitution of split caspases enables selective cell killing. **(A)** An ideal therapeutic caspase would exclusively activate in the presence of a user-defined target protein such as mutant Ras. **(B)** Workflow to screen Ras-targeting caspases (details in Methods). **(C)** Naturally inspired design strategies for engineering mutant Ras-dependent caspases. **(D)** Screening of engineered caspases reveals sensitive and specific designs (upper left corner).

## Results

### Split effector caspases enable mutant Ras-dependent cell killing

Engineering a *retargeted* caspase requires uncoupling it from native activation cues and, instead, making its activation strictly dependent on a chosen target. Human initiator and inflammatory caspases are modular proteins produced as inactive, monomeric zymogens and activated through proximity-induced dimerization, mediated by a distinct recruitment domain (**Figure 1C**, left panel) ^6,13,14^. These steps provide opportunities to restrict caspase activation to diseased cells.

Here, we focus on targeting mutated Ras, an oncogene found in approximately a quarter of all cancer patients ^15–17^, for which targeted inhibitors remain prone to resistance^18–20^. Since mutant Ras forms membrane-localized nanoclusters^15,21^, we hypothesized that initiator and inflammatory caspases could be retargeted to Ras by replacing their recruitment domains with mutant-selective Ras binders, such as the fibronectin-derived human monobody 12VC1^22^, which specifically recognizes KRAS^G12C^ (**Figure 1C**).

To test this design hypothesis, we expressed caspase constructs and Ras in HEK293FT cells and quantified cell death using Annexin V staining (**Figure 1B. Methods**). Contrary to expectations, the engineered caspases produced neither strong killing nor meaningful discrimination between mutant and wild-type Ras (**Figure 1C**, left panel, **1D, Figure S1A**). Other designs based on initiator, as well as inflammatory caspases, also failed (**Figure 1D. Figure S1B-C**). These failures could reflect the dynamic nature of Ras nanoclusters^21^, or the weakness of initiator caspase-caspase interactions prior to self-processing (~µM *K*_*d*_) ^23^, both of which could reduce formation of productive activation complexes.

In contrast to initiator and inflammatory caspases, effector caspases dimerize with nanomolar affinity ^23^, potentially enabling them to better detect transient Ras nanoclusters. However, effector caspases are normally activated by proteolytic cleavage between their subunits by an upstream protease ^6,13,14^, making it unclear how to couple their activity to a non-protease target (**Figure 1C**). To bypass dependence on input proteases, we synthetically split the effector caspase into its large and small subunits and separately fused each to 12VC1. This yielded a split enzyme in which Ras-dependent co-localization could drive caspase reconstitution and activation (**Figure 1C**, right panel).

**Figure S1.**
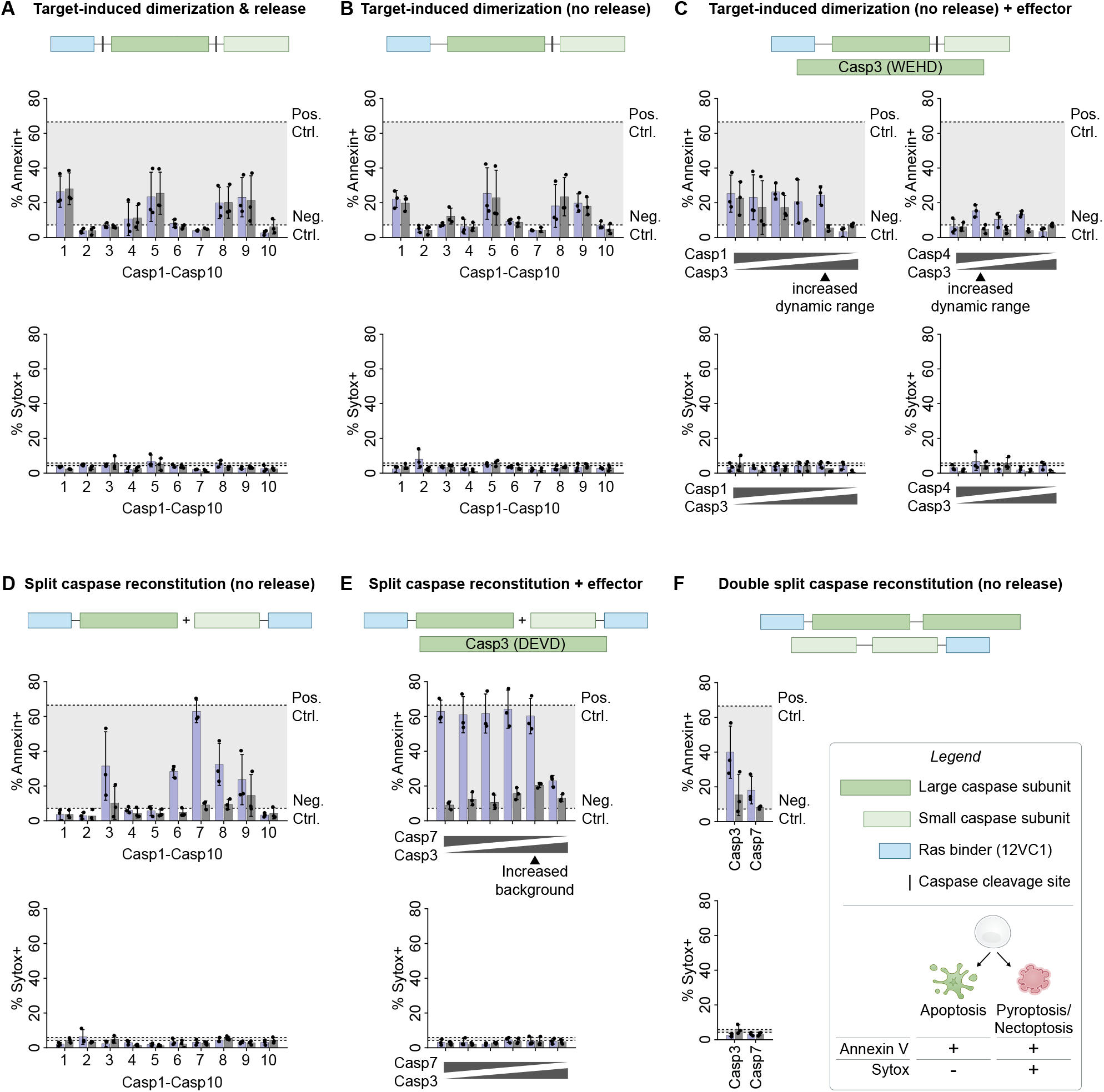
Overview of synthetic caspase designs tested for Ras targeting. **(A)-(F)** Purple and gray bars represent mean cell death (fraction of Annexin- or Sytox-positive cells) for KRAS^G12C^ and KRAS^WT^, respectively. Error bars indicate s.d. across replicates. **(C), (E)** Sensor-caspase plasmid amounts (left to right): 30, 25, 20, 15, 10 (except for Casp4), and 0 ng. Ectopic effector-caspase plasmid amounts (left to right): 0, 5, 10, 15, 20 (except for Casp4), and 30 ng.

Remarkably, split effector caspases enabled sensitive and specific detection of mutant Ras (**Figure 1C**, right panel, **1D, S1D**). All three human effector caspases (caspase-3, -6, and -7) exhibited low background in the presence of wild-type Ras (comparable to NeoR negative control) yet drove robust killing in the presence of ectopic mutant Ras. With caspase-7, this on-target killing approached the efficiency of the constitutively active caspase positive control (**Figure 1C**, right panel). Interestingly, this design was not restricted to effector caspases, as it also enabled Ras-mutant specificity with the initiator caspase-8 (**Figure S1D**). Overall, the split caspase-7 outperformed more than 60 other retargeted caspase designs tested here (**Figure 1D. S1-S2**). We termed this protein *Raspase-7*. Taken together, these data establish a strategy for retargeting caspase activation to mutant Ras.

### Synthetic Raspase-7 recapitulates native caspase activation

Despite being engineered, Raspase-7 is built around the same two principles that govern native caspase regulation: proximity-induced subunit assembly and the separation of recruitment from catalysis^6,13,14^. We therefore asked whether this synthetic enzyme recapitulates features of native caspase activation beyond its cell-killing output.

In their active form, native effector caspases form complexes comprised of two large and two small subunits ^6,13,14^. Raspase-7 killing likewise required both subunits (**Figure 2A**) and scaled with caspase and Ras expression (**Figure 2B. Figure S2A**). This requirement raised an apparent paradox: assembling all four subunits directly through Ras binding seemed unlikely, since Ras nanoclusters are dynamic and co-recruiting four diffusing chains would be inefficient. Yet, reconstituting a single large-small heterodimer should, on its own, be insufficient for activity. To resolve this, we linked the split subunits to complementary coiled-coil heterodimerization domains^24^. This construct restored caspase activity (**Figure S2B**), suggesting that reconstituted split proteases self-assemble into active caspase complexes (**Figure S2C**). These results suggest a model in which Ras-dependent co-localization promotes reconstitution of the large and small subunits, which subsequently homodimerize through intrinsic caspase-caspase interactions to form active complexes. This model draws on features of both caspase classes^6,13,14^. As with initiators, caspase-caspase interactions are gated by recruitment. As with effectors, interactions are favored by the high intrinsic affinity between subunits.

**Figure 2.**
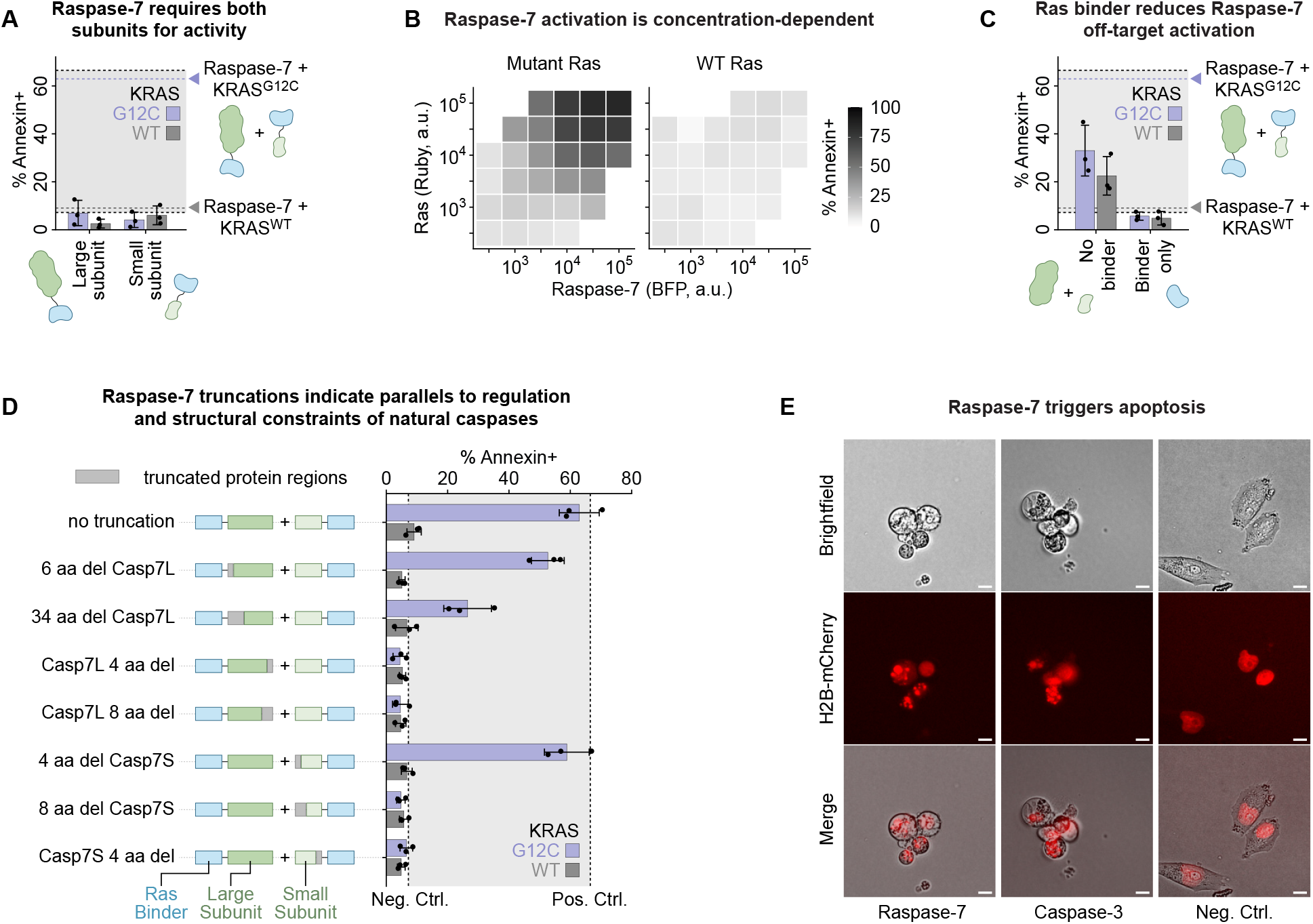
Raspase-7 mechanism mirrors features of natural initiator and effector caspases. **(A)** Raspase-7 requires both subunits for activity. **(B)** Raspase-7 activation is concentration-dependent. **(C)** Raspase-7 requires fusion to a Ras-binding domain for background suppression. **(D)** Truncation studies show that Raspase-7 recapitulates key mechanistic features of native caspase activation. **(E)** Raspase-7 triggers apoptotic cell death (membrane blebbing, nuclear fragmentation) in Ras-mutant cells (MIA PaCa-2, co-transfection of H2B-mCherry). Scale bar: 10 µm.

A hallmark of native caspases is their low activity in the absence of a death trigger. Notably, Raspase-7 retained a low background even when overexpressed. To understand why its subunits do not spontaneously reconstitute, we compared constructs with and without the 12VC1 Ras-binding domain. Unlike full-length Raspase-7, control constructs lacking 12VC1 induced death in a Ras-independent manner (**Figure 2C**), indicating that the 12VC1 domain actively suppresses spontaneous subunit assembly, functionally analogous to the autoinhibition imposed by recruitment domains in initiator caspases^14^.

**Figure S2.**
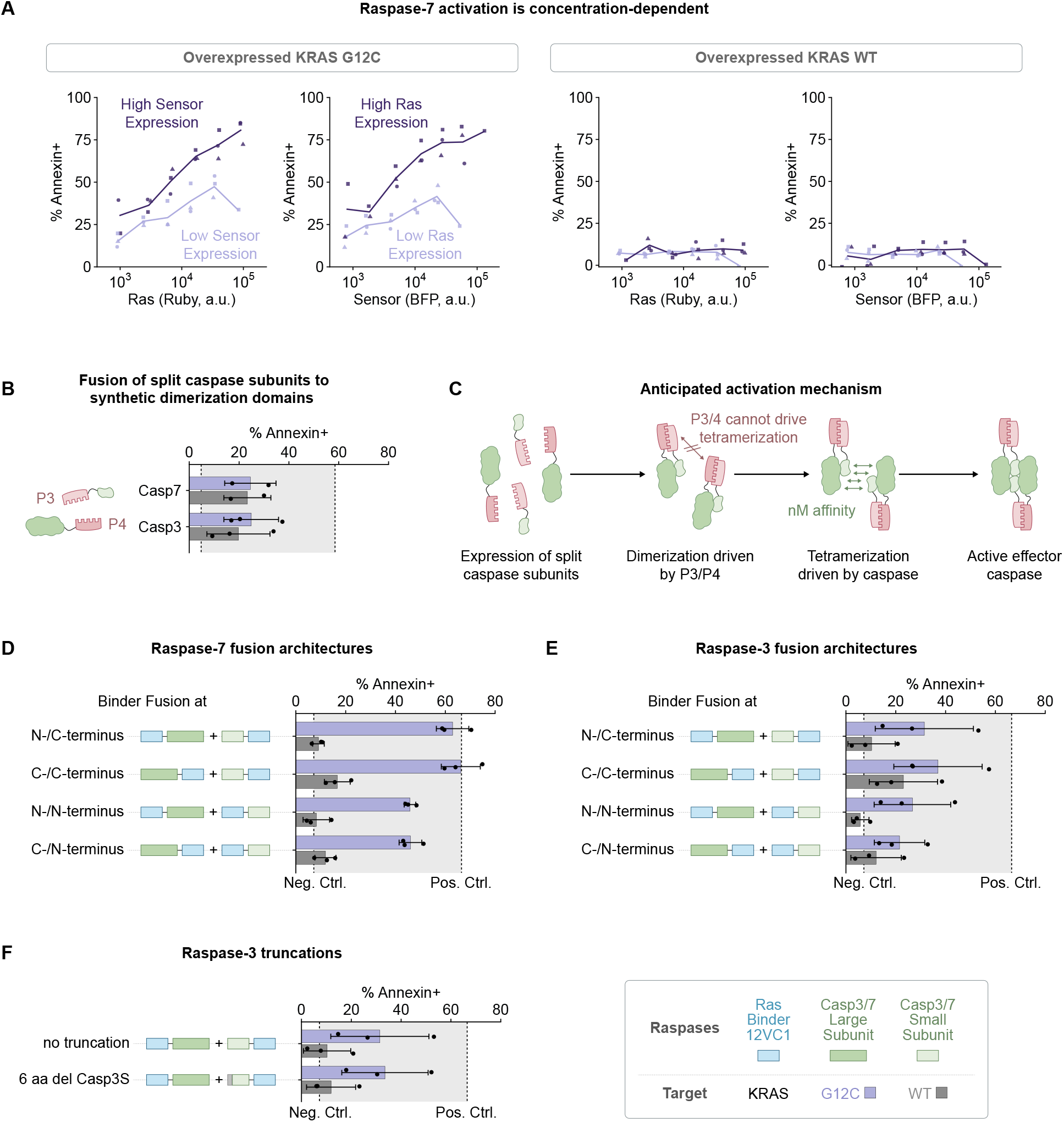
Mechanistic characterization of Raspase-7. **(A)** Raspase-7 activation depends on both Ras abundance and circuit expression. Ruby and BFP fluorescence were used to quantify Ras and caspase-sensor expression, respectively. Data points were generated by binning Ruby and BFP signals into discrete expression ranges; marker shapes are matched across replicates. **(B)** Cell death induced by constructs in which the P3/P4 dimerization domains were fused to complementary caspase subunits. Flow-cytometry gating: Ruby > 10^3^ and 10^4^ < BFP < 10^4.5^. Purple and gray bars report mean cell death (fraction Annexin V-positive cells) for KRAS^G12C^ and KRAS^WT^, respectively; error bars denote s.d. across replicates. **(C)** Proposed activation mechanism of split caspases: Dimerization of split caspase subunits is sufficient for sensor activation and cell death induction. **(D)-(F)** Raspases recapitulate key mechanistic features of native caspase activation: Purple and gray bars represent mean cell death for KRAS^G12C^ and KRAS^WT^, respectively; error bars indicate s.d. across replicates. **(D), (E)** Comparison of fusion architectures for Raspase-7 **(D)** and Raspase-3 **(E). (F)** Evaluation of truncated circuit variants for Raspase-3.

Also consistent with native caspase regulation, *N*-terminal fusion of 12VC1 to the small subunit reduced maximal activation relative to *C*-terminal fusion (**Figure S2D**), in line with the known contribution of the small-subunit *N*-terminus to active-site geometry^25,26^. Truncation studies (**Figure 2D**) and analogous experiments with the split caspase-3 design (“*Raspase-3*”, **Figure S2E, F**) further supported parallels to endogenous regulation ^6,25–28^.

Finally, Raspase-7 killing displayed key hallmarks of apoptosis such as membrane blebbing and nuclear fragmentation ^29^ (**Figure 2E. Figure S1D**), confirming that it acts as a *bona fide* effector caspase. Together, these results indicate that retargeted split caspases preserve key constraints of native activation, while leveraging target proximity and fusion-imposed constraints to achieve specificity.

### Raspases selectively eliminate Ras-mutant cancer cells

To function therapeutically, a retargeted caspase should (i) be amenable to repeated delivery via a clinically relevant and safe delivery vehicle, (ii) respond to disease markers at physiological levels, and (iii) induce potent cell killing. It should also incorporate desirable features for interaction with the immune system, including (iv) avoiding non-human sequences to reduce the risk of undesired immunogenicity and (v) enabling controlled recruitment of immune cells, for example, through the induction of the pyroptosis cell death program^7,30,31^, which facilitates elimination of non-transfected tumor cells (**Figure 3A**).

**Figure 3.**
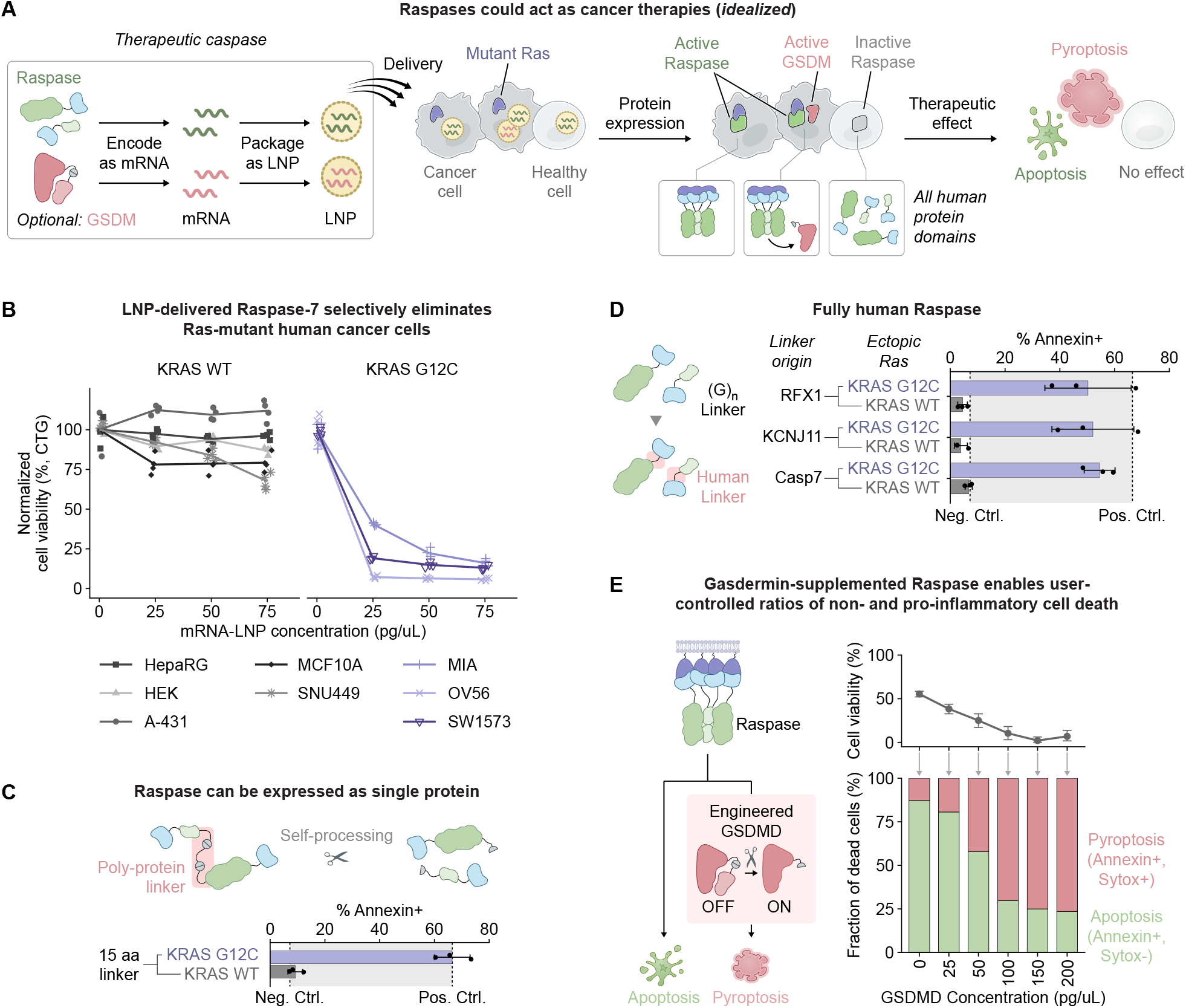
LNP-delivered Raspases potently and selectively eliminate Ras-mutant cancer cells. **(A)** Raspases could act as cancer therapies (*idealized*). **(B)** Cell viability of wild-type and mutant Ras human cancer cell lines 3 days (3d) after mRNA-LNP Raspase-7 transfection. **(C)** Raspase-7 functions when expressed as a single self-cleaving polyprotein. **(D)** Raspase-7 functions with human protein linker sequences. **(E)** Co-delivery of engineered GSDMD enables modulation of relative rates of apoptosis and pyroptosis.

**Figure S3.**
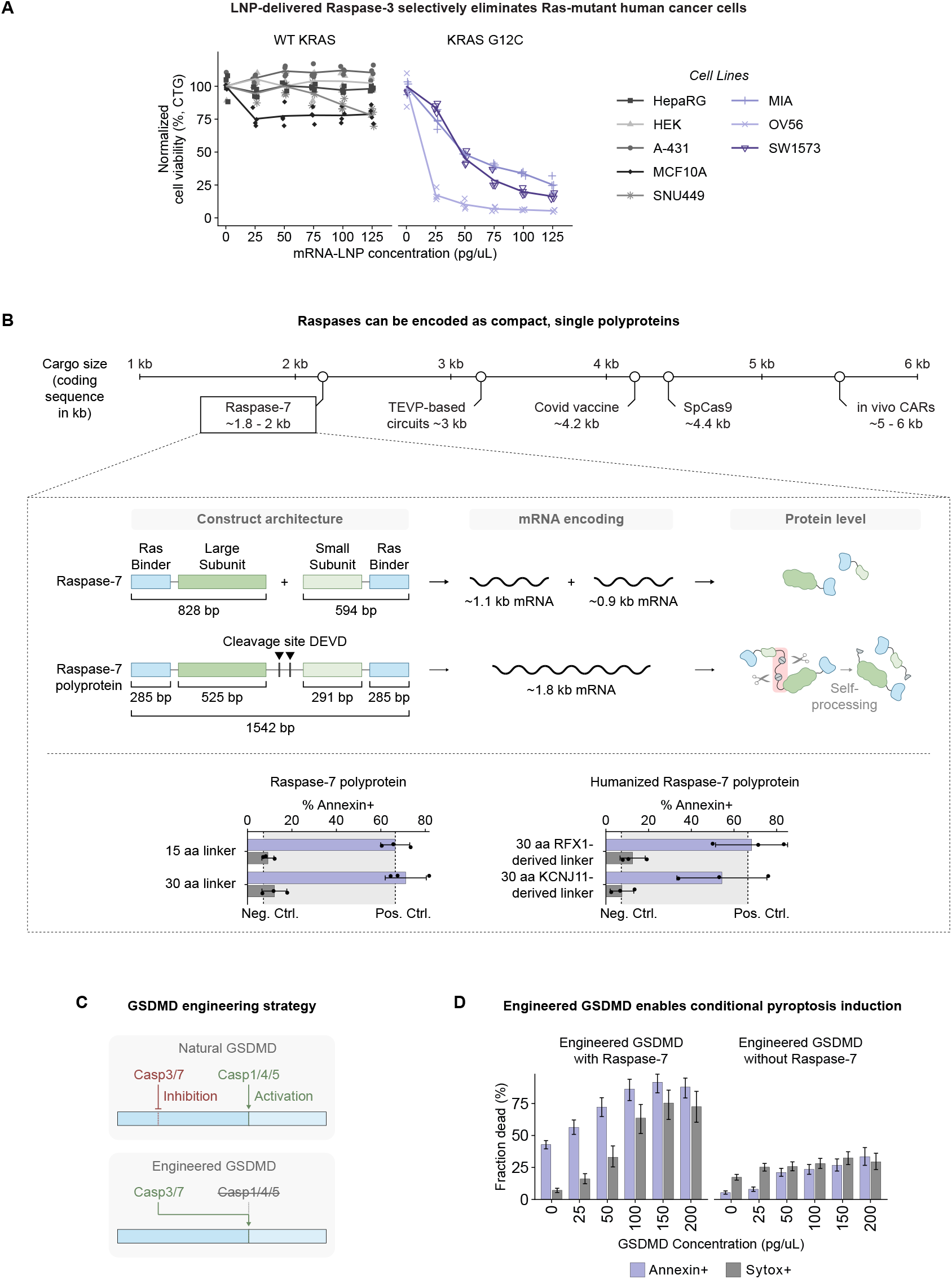
Raspases could have therapeutic utility. **(A)** LNP-delivered Raspase-3 selectively eliminates Ras-mutant cancer cells. Cell viability was measured 3d after circuit treatment by CellTiter-Glo and normalized to a mock LNP transfection control encoding NeoR. Points denote replicates; lines indicate the replicate mean. **(B)** Raspases have a compact cargo footprint that can be further reduced by expressing the circuit as a single polyprotein. Self-processing is the anticipated mechanism, but was not experimentally validated here. Purple and gray bars report mean cell death for KRAS^G12C^ and KRAS^WT^, respectively; error bars denote s.d. across replicates. **(C)** Raspases can be coupled to gasdermins to induce pyroptosis: Schematic of the GSDMD engineering strategy. **(D)** Fraction of Annexin V-positive and Sytox-positive cells following delivery of engineered GSDMD-supplemented Raspase-7. MIA PaCa-2 cells received either 150 pg/µL sensor (left) or 150 pg/µL NeoR (right) formulated as mRNA-LNP, together with 150 pg/µL H2B-mCherry mRNA as a co-transfection marker. Flow-cytometry gating: mCherry > 10^3^. Bars denote means; error bars indicate bootstrap-derived 95% confidence intervals.

To evaluate deliverability, target sensitivity, and killing potency, we encoded Raspases as mRNAs, encapsulated them in lipid nanoparticles (LNPs), and treated human cancer cell lines harboring endogenous wild-type or mutant Ras (**Methods**). As measured by CellTiter-Glo, Raspase-7 selectively eliminated Ras-mutant lines, while sparing Ras-wild-type lines (**Figure 3B**). Raspase-3, the analogous design based on effector caspase-3, was less sensitive than Raspase-7 but still achieved near-complete elimination at higher doses (**Figure S3A**), supporting the generality of the split-effector design. Together, these results demonstrate mRNA-LNP deliverability and potent but selective cytotoxicity.

To compress the multi-protein Raspase system described above into a single protein, inspired by self-cleaving proteases in viral polyproteins ^32^, we re-engineered Raspase-7 as a polyprotein whose concatenated protein components were separated by self-cleavage sites (**Figure 3C**). This compression decreased mRNA payload from ~2 kb to ~1.8 kb, while maintaining potency and specificity (**Figure S3B**).

To reduce the risk of undesired immunogenicity arising from Raspase expression in healthy cells, we also sought to eliminate the non-human sequences that had been incorporated in various linkers. Replacing synthetic linkers with flexible, glycine-rich segments derived from the human proteins RFX1, KCNJ11, and CASP7 preserved efficacy compared to the original designs (**Figure 3D. Figure S3B**).

Therapeutic applications of Raspases could benefit from selective induction of pro-inflammatory pyroptotic cell death in cancer cells, as pyroptosis can promote immune-mediated elimination of neighboring non-transfected tumor cells^7,30,31^. Pyroptosis is executed by gasdermins, pore-forming proteins naturally activated by caspase cleavage^7,33^. We reasoned that an engineered gasdermin activatable by Raspase but not by endogenous inflammatory caspases could convert Raspase output from apoptosis to pyroptosis. We therefore re-engineered GSDMD, one of the most potent gasdermins, by replacing its activating cleavage site and removing inhibitory cleavage sites^34^, to create a pyroptotic “*diverter*” (**Figure S3C**). When co-delivered with Raspase-7, the diverter triggered rapid pyroptosis, as confirmed by Sytox staining (**Figure S3D. Methods**). By contrast, when delivered alone, it exhibited much lower cell death. Further, varying the relative concentration of diverter and Raspase enabled systematic modulation of the relative frequencies of apoptosis and pyroptosis (**Figure 3E**), potentially allowing tuning of immune responses.

### Raspase-7 outperforms other RAS-targeted interventions

Finally, we compared the Raspase construct to alternative Ras-targeting interventions. Clinically, intracellular Ras is primarily targeted with small-molecule inhibitors, including mutation-selective agents (*e*.*g*., Sotorasib^35,36^) and emerging pan-Ras inhibitors (*e*.*g*., RMC-7977, the clinical tool compound for Daraxonrasib^16,37,38^). As an additional comparator, we included recent Ras-targeting protein circuits ^39^. These circuits use a multi-step mechanism in which mutant Ras nanoclusters drive reconstitution of the viral split protease TEVP (Tobacco Etch Virus Protease), which, in turn, proteolytically activates caspase-3.

When delivered at equal mRNA dose, Raspase-3 performed comparable to or better than TEVP-based circuits, depending on the cell line. Raspase-7 showed higher sensitivity than TEVP circuits across all cell lines, with only a modest increase in Ras-wild-type background (**Figure 4A. Figure S4**). Further, both retargeted caspase sensors outperformed the pharmacological Ras inhibitors RMC-7977 and Sotorasib (data from Lu *et al*. ^*39*^). Notably, two of the three tested cancer cell lines were not addicted to Ras and did not respond to the inhibitors, highlighting the value of sense-and-kill strategies over oncogene inhibition.

**Figure 4.**
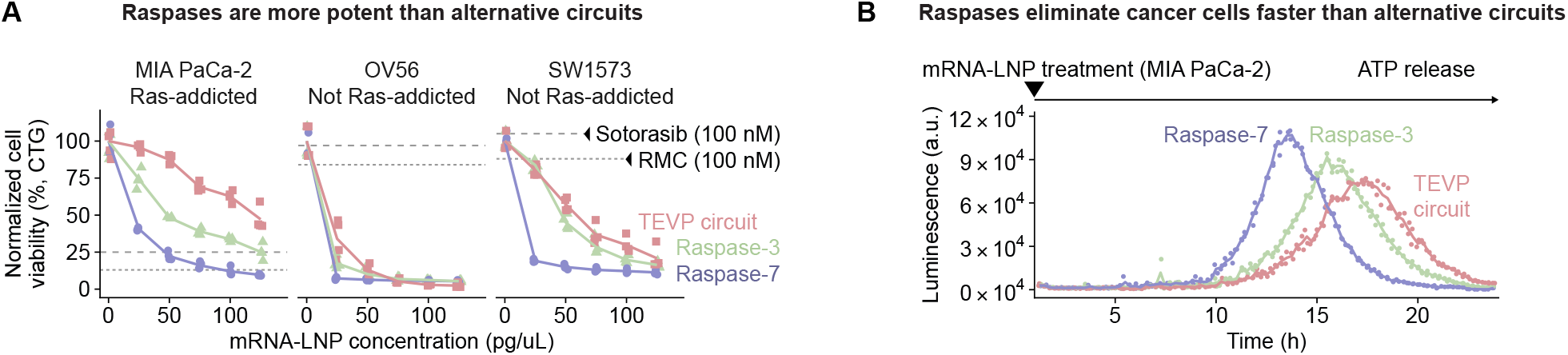
Raspase-7 has benefits compared to alternative Ras-targeting interventions. **(A)** Viability of mutant Ras human cancer cell lines 3d after treatment. Drug benchmarks based on data in Lu *et al* ^*39*^. **(B)** Raspase-7 eliminates Ras-mutant MIA PaCa-2 cells more rapidly than TEVP-based circuits.

We also compared the kinetics of cell death execution: As determined by ATP release measurements, Raspase-7 killed Ras-mutant cancer cells 1.3-fold faster than TEVP circuits (**Figure 4B. Methods**). Thus, despite their smaller size, single-step mechanism, and restriction to human sequences, the Raspase proteins were able to provide specificity and potency comparable to, or better than, those of existing drugs and circuits.

**Figure S4.**
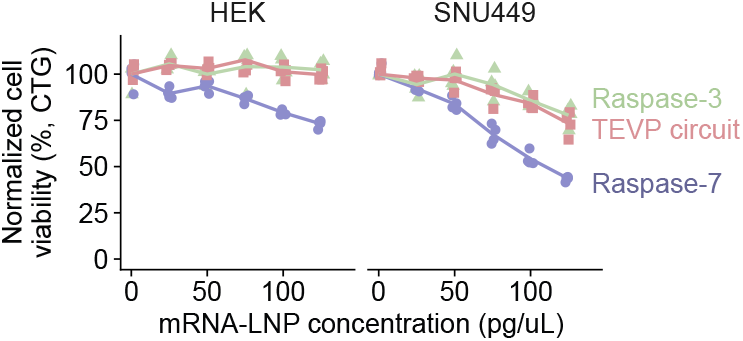
Comparison of Raspases and TEV protease-based Ras-targeting circuits in wild-type Ras cells. Viability of KRAS^WT^ cells was measured by CellTiter-Glo 3d after treatment and normalized to the NeoR mock transfection control. Points denote replicates; lines indicate the replicate mean.

## Discussion

Caspases are the cell’s most direct executioners of programmed cell death ^1^. However, their therapeutic utility has been limited by the difficulty of coupling their activity to disease-specific inputs. By drawing on two universal features of natural caspase regulation—proximity-induced subunit assembly and the modular separation of recruitment from catalysis—we designed a synthetic caspase, *Raspase*, that selectively eliminates Ras-mutant cells.

The Raspase proteins recapitulate many of the hallmarks of native caspase biology while operating under entirely new control. They remain inactive as separated subunits and assemble into active complexes only upon detection of their target, much like endogenous initiator zymogens^6,13,14^. They also preserve key regulatory constraints of native effector caspases and execute authentic death programs consistent with hallmark signatures of apoptosis^14,25–29^. Finally, supplemented engineered gasdermins can divert their output from non-inflammatory apoptosis to pro-inflammatory pyroptosis, mirroring native pyroptosis induction cascades^7,33^.

Raspases define a therapeutic strategy distinct from existing approaches. Pan-caspase activators or inducible suicide switches such as iCasp9 trigger apoptosis efficiently but are not selective for diseased cells^9,10,40,41^. Targeted therapies, including Ras inhibitors, achieve selectivity but act indirectly by perturbing endogenous signaling, often inducing proliferative arrest rather than active killing ^16,18–20,39^. Other synthetic sense-and-kill circuits have been described, but typically do not directly retarget human caspases, require multiple components, rely on foreign protein components, or face delivery constraints, any of which may complicate translation ^39,42–47^. Raspases unite the direct lethality of a suicide switch with the selectivity of a targeted inhibitor into a single, mRNA-encoded human polyprotein deliverable by LNPs. Notably, Raspases efficiently eliminated cancer lines not addicted to RAS that resisted pharmacological inhibition, highlighting the value of sense-and-kill strategies that couple detection directly to death over approaches that function indirectly by inhibiting oncogene function.

Finally, Raspases are built from interchangeable sensing and catalytic modules, making them inherently reprogrammable. Substituting the Ras-binding monobody for another intracellular binder—a monobody, nanobody, or other human-derived domain—should redirect killing to new targets without re-engineering the catalytic core. Combining binders could enable AND-gated recognition of marker combinations to sharpen specificity. Conversely, co-delivering Raspases recognizing multiple targets could expand specificity through OR logic. More broadly, our results establish retargeted caspases as a reprogrammable sense-and-respond platform for cancer and other diseases that require selective elimination of diseased cells.

### Limitations of the study

Although Raspase-7 and engineered GSDMD constructs exhibit low background, residual target-independent killing emerges at high doses, and the affinities, assembly kinetics, and stoichiometry of the reconstituted Raspase complexes remain incompletely characterized. Our validation used cancer cell lines with defined Ras status and did not capture the full heterogeneity of mutant and wild-type Ras levels clinically observed across patient tumors, nor the range of Ras mutations beyond those recognized by the 12VC1 monobody. The generality of the platform to other intracellular targets, while conceptually straightforward, was not tested here and will depend on the availability of selective binders, which will benefit from recent improvements in computational binder design^48–51^. Finally, the immunogenicity of the engineered proteins, though minimized by restricting sequences to human domains, was not directly measured. Moreover, junctions between fused domains necessarily introduce non-native sequences that may harbor immunogenic epitopes, a limitation addressable by established *in silico* and experimental methods ^52–55^.

## Methods

### Plasmid construction

Plasmids were assembled using Gibson Assembly (NEBuilder HiFi DNA Assembly Master Mix; New England BioLabs) or KLD cloning (T4 Polynucleotide Kinase, T4 DNA Ligase, DpnI, T4 DNA Ligase Buffer; Thermo Scientific). Gene fragments were obtained from Twist Bioscience, Integrated DNA Technologies (IDT), or generated by PCR amplification from existing constructs. All PCR primers were synthesized by IDT. Prior to experiments, plasmids were purified (QIAprep Spin Miniprep Kit or Qiacube machine; Qiagen), concentration-normalized (Thermo Scientific NanoDrop 8000), and sequence-validated (Plasmidsaurus, Genewiz, or Quintara).

### Tissue culture and cell lines

Cell lines were sourced from ATCC and maintained under standard conditions (37°C, 5% CO_2_, humidified Eppendorf CellXpert C170i incubator). We used Dulbecco’s Modified Eagle Media (DMEM; Thermo Fisher) supplemented with 10% fetal bovine serum (FBS; Avantor), penicillin (1 unit/ml), streptomycin (1 µg/ml), glutamine (2 mM), sodium pyruvate (1 mM), and 1X Minimal Essential Media Non-Essential Amino Acids (all Thermo Fisher) to culture cells. HEK293FT cells were used for up to 20 passages, other human cancer cell lines for up to 10 passages, with cultures maintained at 10%-90% confluency. Cells were passaged using 0.25% Trypsin-EDTA (Thermo Fisher). We used Trypan Blue (Invitrogen) and the Countess 3 automated cell counter (Thermo Fisher) to determine viable cell counts and calculate seeding densities. Mycoplasma status was routinely checked using the MycoStrip kit (InvivoGen).

### DNA transfection for caspase screening

In order to screen caspase designs, we expressed caspase constructs in HEK293FT cells together with mutant (KRAS^G12C^) or wild-type (KRAS^WT^) Ras, and quantified apoptosis 20-24 h later by Annexin V and Sytox staining (see *flow cytometry for caspase screening*). FuGENE HD (Promega) was used for transient DNA transfections. Unless noted otherwise, we plated 50k HEK293FT cells per well in 96-well plates and performed reverse transfections immediately after seeding, following the manufacturer’s instructions. More specifically, we poly-transfected cells with 5 ng of Ras (35 ng co-transfected mRuby3) and 30 ng of circuit DNA (30 ng co-transfected mTagBFP2). We used 6 µL of transfection reagent per 1 µg of plasmid DNA. For split circuits, we transfected 15 ng of each complementary plasmid unless otherwise noted. We used a TEVP-activated caspase-7 as a positive control, inspired by the design of Xia *et al*. ^*44*^, and NeoR as a negative control in all DNA transfection experiments unless otherwise noted. We followed the same experimental procedure for validation of single-protein Raspases as well as Raspases with human linker sequences.

### Flow cytometry for caspase screening

Cells were harvested between 20-24h post-transfection for flow cytometry. To enable analysis of all cells, independent of their viability, we first collected the supernatant with detached cells. Remaining cells were trypsinized and pooled with cells in the supernatant. We then added an excess of Hank’s Balanced Salt Solution (HBSS, Gibco) containing 2.5 mg/ml bovine serum albumin (BSA), 2.5 mM calcium chloride, 15 µL/mL Sytox Green Ready Flow reagent (R37168, Invitrogen), and 15 µL/mL Annexin V Ready Flow reagent (R37176, Invitrogen). After 5 minutes of incubation, cells were passed through a 40-µm cell strainer to remove aggregates and analyzed with a CytoFLEX S flow cytometer (Beckman Coulter). Unless otherwise specified, fluorescence was measured using FITC-A (Sytox; excitation 488 nm, emission 525/40 nm; gain 1), ECD-A (mRuby3 or mCherry; excitation 561 nm, emission 610/20 nm; gain 1), PB450-A (mTagBFP2; excitation 405 nm, emission 450/45 nm; gain 1), and APC-A (Annexin V; excitation 638 nm, emission 712/25; gain 1). Gating to remove artifacts or cell aggregates based on forward- and side-scatter was performed in FlowJo (version 10.10, BD Biosciences), without removing dead cells. Gates for fluorescent protein expression were applied in Python as specified in **Figure 1B**. Briefly, for caspase screening experiments, we used the following screening window: 10^4^ < Ruby (Ras) < 10^4.5^ and 10^4^ < BFP (circuit) < 10^4.5^. Across all experiments, cells with Annexin V signal larger than 10^3.1^ were counted as Annexin-positive. All cells with Sytox signal larger than 10^3.2^ were denoted as Sytox-positive. We followed the same experimental procedure as outlined here for validation of single-protein Raspases as well as Raspases with human linker sequences.

### Cell imaging

We imaged live and dead cells on an EVOS FL Auto cell imaging system (Life Technologies) ~16-24 hours after mRNA-LNP transfection using an EVOS RFP (Ex 542/20 nm, Em 593/40 nm) filter cube. Images were saved as TIFF files and processed using a custom Python script. Within each figure panel, images were displayed using identical brightness and color scaling.

### mRNA production

Templates for transcription were generated by PCR linearization (Q5 High-Fidelity DNA Polymerase, NEB) of DNA constructs containing a 5’ T7 promoter followed by the dinucleotide AG. A 120-base pair polyA tail was introduced at the 3’ end by PCR. Linear DNA was purified using gel extraction (Zymoclean Gel DNA Recovery kit, Zymo) or PCR purification (DNA Clean & Concentrator kit, Zymo). mRNA was then synthesized by *in vitro* transcription using NEB’s HiScribe T7 High Yield RNA Synthesis Kit. The IVT mixture consisted of 10X Reaction Buffer (1X final, NEB), 1 µg DNA template, 4 mM CleanCap AG (TriLink), 2 µl T7 RNA polymerase mix (NEB), and ATP, GTP, CTP (all NEB), and *N1*-Methyl-Pseudouridine-5’-Triphosphate (TriLink) each at 5 mM final concentration. Reactions were incubated for 2 hours at 37 °C, followed by the addition of DNAse I (NEB) and a further 15-minute incubation at 37 °C. Purification was performed using Zymo’s RNA Clean and Concentrator kit. mRNA concentrations were measured by Nanodrop or Qubit (RNA High Sensitivity or Broad Range kits, Invitrogen). Concentration-normalized mRNAs were stored at −80 °C.

### LNP production

We used a previously reported 4-component LNP formulation (4A3-SC8, DOPE, Cholesterol, DMG-PEG) for all LNP transfections^56,57^. We followed instructions in Wang *et al*. for the preparation of the 4-component lipid mastermix ^57^. For encapsulation, the working lipid mixture was equilibrated at room temperature for at least 5 minutes, then vortexed for 5 seconds. A lipid mix was then prepared by combining 12 µL lipid mastermix with 18 µL 200-proof ethanol (30 µL per reaction) and aliquoted into separate tubes. RNA mixtures were prepared by combining 40 µL RNA (250 ng/µL; 10 µg total input) with 32 µL nuclease-free water and 18 µL 50 mM citrate buffer, for a total of 90 µL per reaction. LNPs were formed by vortexing 30 µL lipid mastermix and rapidly dispensing the 90 µL RNA mixture into the lipid phase; mixing was continued for 30 seconds. After mixing, the dispersions were held at room temperature for 5 minutes, and dialysis was started within 15 minutes. Pur-A-Lyzer Midi 3500 dialysis tubes were used and preconditioned by filling with 900 µL water for 5 minutes. Approximately 120 µL of each LNP sample was loaded into a tube, placed in a styrofoam holder, and submerged in PBS in a beaker. Dialysis proceeded either for 1 hour at room temperature or overnight at 4 °C. Post-dialysis, samples were transferred into RNase-free 1.5 mL microcentrifuge tubes, and the final volumes were recorded. Each sample was adjusted to 500 µL with PBS and stored at 4 °C. 10% sucrose was added to enable LNP storage at −20 °C.

### mRNA-LNP transfection of human cancer cell lines

LNPs were reverse-transfected by first adding the corresponding dose of LNPs to 96-well culture plates, then adding 2k cells on top to a total volume of 100 µL per well. For split caspases, the same mRNA amounts of both split enzyme halves were co-encapsulated into a single LNP. Nunc Edge 2.0 plates were used to reduce edge effects. Unless otherwise specified, total mRNA-LNP dose was normalized to 300 pg/µL by supplementing formulations with NeoR mRNA-LNP as a matched filler.

### Cell viability assay with CellTiter-Glo

Cell viability was measured with the CellTiter-Glo (CTG) 2.0 Luminescent Cell Viability Assay (Promega). On the assay day (3 days after transfection unless otherwise specified), plates were brought to room temperature for 30 min before adding CTG. CTG reagent (100 µL) was added directly to each well, and plates were shaken for 2 min on an orbital shaker within a Promega GloMax instrument to promote lysis and mixing. The plates were then incubated at room temperature for 10 min to allow the luminescent signal to stabilize. After incubation, 180 µL from each well was transferred to a white plate suitable for luminescence detection, and luminescence was measured on the GloMax according to the manufacturer’s instructions.

### Cell viability assessment for pyroptosis-inducing Raspases

Since the CellTiter-Glo (CTG) assay cannot distinguish between apoptosis and pyroptosis, we quantified viability by flow cytometry in experiments that use both caspases and gasdermins. Briefly, 10k MIA PaCa-2 cells were reverse-transfected with 150 pg/µL of caspase sensor mRNA-LNP and the amount of gasdermin mRNA-LNP as annotated in figures. Further, constructs were co-transfected with 150 pg/µL H2B-Cherry, enabling gating for transfected cells (mCherry > 10^3^). The total amount of LNP was filled up to 500 pg/µL using a NeoR mRNA-LNP (negative control) to reduce effects from differences in LNP transfection. Flow cytometry was performed as described above 16-20h after transfection.

### ATP release assay

Briefly, 5k MIA PaCa-2 cells per well were plated in white-bottom 96-well plates in DMEM. Approximately 4h after seeding, treatments were applied (200 pg/µL), and RealTime-Glo Extracellular ATP Assay Reagent (Promega) was added directly to each well. After orbital shaking to mix, extracellular ATP (ATP release) was monitored over 24 hours (outside of a CO_2_-controlled environment).

### Drug treatment

Treatment concentrations of 100 nM are shown. Sotorasib (MedChemExpress, Cat. No.: HY-114277) and RMC-7977 (MedChemExpress, Cat. No.: HY-156498) efficacy was calculated from data in Lu *et al*. ^*39*^

### Software

The following software was used in this study: Cursor, FlowJo, Python, R, AlphaFold 3.

### Statistics and reproducibility

All experiments were performed with at least three technical and/or biological replicates unless otherwise noted.

## Resource availability

### Data Availability

Upon final publication, all raw data, including ungated flow cytometry data and raw microscopy images, will be deposited at *data*.*caltech*.*edu* and made publicly available. Upon final publication, key plasmids will be made available via Addgene. Requests for materials and correspondence should be addressed to M.B.E.

### Code Availability

Upon final publication, all code used for data analysis and figure generation will be made publicly available at *data*.*caltech*.*edu*. Any additional information required to reanalyze the data reported in this paper is available from the lead contact upon request.

## Acknowledgements

We thank Shiyu Xia, James Linton, Dongyang Li, Nikita Makarov, and Albert Qiang for advice on caspase engineering and experimental design, insightful discussions, and critical proofreading of the manuscript; Inna-Marie Strazhnik for graphical design; Leah Santat and Rui Malinowski for administrative support. This research was supported by the National Institutes of Health (*EB030015*), the Mathers Foundation (*12540447*), the Yosemite-American Cancer Society Award, and the Curci Foundation (Caltech Cell Interaction Initiative). The content is solely the responsibility of the authors and does not necessarily represent the official views of the National Institutes of Health. M.B.E. is a Howard Hughes Medical Institute Investigator. L.M. and A.C.L. are supported by the Merkin Institute for Translational Research (Merkin Bridge Fellowship).

## Author contributions

L.M., A.C.L., and M.B.E. conceived and designed the study. M.B.E. directed and supervised the study. L.M., A.C.L., K.H., and E.Z. performed all experiments. L.M. and A.C.L. analyzed data. L.M., A.C.L., and M.B.E. wrote the manuscript with input from all authors.

## Declaration of generative AI in the manuscript preparation process

During the preparation of this work, the authors used Claude and ChatGPT to improve the language and readability of the manuscript text. After using these tools, the authors reviewed and edited the content as needed and take full responsibility for the content of the published article.

## References

1. Fuchs, Y. & Steller, H. Programmed cell death in animal development and disease. Cell 147, 742–758 (2011).

2. Taylor, R. C., Cullen, S. P. & Martin, S. J. Apoptosis: controlled demolition at the cellular level. Nat. Rev. Mol. Cell Biol. 9, 231–241 (2008).

3. Arandjelovic, S. & Ravichandran, K. S. Phagocytosis of apoptotic cells in homeostasis. Nat. Immunol. 16, 907–917 (2015).

4. Morioka, S., Maueröder, C. & Ravichandran, K. S. Living on the edge: Efferocytosis at the interface of homeostasis and pathology. Immunity 50, 1149–1162 (2019).

5. Thornberry, N. A. & Lazebnik, Y. Caspases: enemies within. Science 281, 1312–1316 (1998).

6. Riedl, S. J. & Shi, Y. Molecular mechanisms of caspase regulation during apoptosis. Nat. Rev. Mol. Cell Biol. 5, 897–907 (2004).

7. Liu, X., Xia, S., Zhang, Z., Wu, H. & Lieberman, J. Channelling inflammation: gasdermins in physiology and disease. Nat. Rev. Drug Discov. 20, 384–405 (2021).

8. Dhani, S., Zhao, Y. & Zhivotovsky, B. A long way to go: caspase inhibitors in clinical use. Cell Death Dis. 12, 949 (2021).

9. Straathof, K. C. et al. An inducible caspase 9 safety switch for t-cell therapy. Blood 105, 4247–4254 (2005).

10. Di Stasi, A. et al. Inducible apoptosis as a safety switch for adoptive cell therapy. N. Engl. J. Med. 365, 1673–1683 (2011).

11. Van Opdenbosch, N. & Lamkanfi, M. Caspases in cell death, inflammation, and disease. Immunity 50, 1352–1364 (2019).

12. Poreba, M., Strózyk, A., Salvesen, G. S. & Drag, M. Caspase substrates and inhibitors. Cold Spring Harb. Perspect. Biol. 5, a008680 (2013).

13. Pop, C. & Salvesen, G. S. Human caspases: activation, specificity, and regulation. J. Biol. Chem. 284, 21777–21781 (2009).

14. Boatright, K. M. et al. A unified model for apical caspase activation. Mol. Cell 11, 529–541 (2003).

15. Simanshu, D. K., Nissley, D. V. & McCormick, F. RAS proteins and their regulators in human disease. Cell 170, 17–33 (2017).

16. Wasko, U. N. et al. Tumour-selective activity of RAS-GTP inhibition in pancreatic cancer. Nature 629, 927–936 (2024).

17. Cerami, E. et al. The cbio cancer genomics portal: an open platform for exploring multidimensional cancer genomics data. Cancer Discov. 2, 401–404 (2012).

18. Ebright, R. Y., Dilly, J., Shaw, A. T. & Aguirre, A. J. Response and resistance to RAS inhibition in cancer. Cancer Discov. OF1–OF25 (2025).

19. Riedl, J. M. et al. Genomic landscape of clinically acquired resistance alterations in patients treated with KRASG12C inhibitors. Ann. Oncol. 36, 682–692 (2025).

20. Sang, B. et al. Mechanisms of resistance to active state selective tri-complex RAS inhibitors (2025).

21. Zhou, Y. & Hancock, J. F. Ras nanoclusters: Versatile lipid-based signaling platforms. Biochim. Biophys. Acta 1853, 841–849 (2015).

22. Teng, K. W. et al. Selective and noncovalent targeting of RAS mutants for inhibition and degradation. Nat. Commun. 12, 2656 (2021).

23. Donepudi, M., Mac Sweeney, A., Briand, C. & Grütter, M. G. Insights into the regulatory mechanism for caspase-8 activation. Mol. Cell 11, 543–549 (2003).

24. Gradišar, H. & Jerala, R. De novo design of orthogonal peptide pairs forming parallel coiled-coil heterodimers. J. Pept. Sci. 17, 100–106 (2011).

25. Chai, J. et al. Crystal structure of a procaspase-7 zymogen: mechanisms of activation and substrate binding. Cell 107, 399–407 (2001).

26. Witkowski, W. A. & Hardy, J. A. L2’ loop is critical for caspase-7 active site formation: Essentiality of l2 loop in caspase-7. Protein Sci. 18, 1459–1468 (2009).

27. Roy, S. et al. Maintenance of caspase-3 proenzyme dormancy by an intrinsic “safety catch” regulatory tripeptide. Proc. Natl. Acad. Sci. U. S. A. 98, 6132–6137 (2001).

28. Thomsen, N. D., Koerber, J. T. & Wells, J. A. Structural snapshots reveal distinct mechanisms of procaspase-3 and -7 activation. Proc. Natl. Acad. Sci. U. S. A. 110, 8477–8482 (2013).

29. Häcker, G. The morphology of apoptosis. Cell Tissue Res. 301, 5–17 (2000).

30. Wang, Q. et al. A bioorthogonal system reveals antitumour immune function of pyroptosis. Nature 579, 421–426 (2020).

31. Zhang, Z. et al. Gasdermin E suppresses tumour growth by activating anti-tumour immunity. Nature 579, 415–420 (2020).

32. Kräusslich, H. G. & Wimmer, E. Viral proteinases. Annu. Rev. Biochem. 57, 701–754 (1988).

33. Broz, P. Pyroptosis: molecular mechanisms and roles in disease. Cell Res. 35, 334–344 (2025).

34. Exconde, P. M., Bourne, C. M., Kulkarni, M., Discher, B. M. & Taabazuing, C. Y. Inflammatory caspase substrate specificities. MBio 15, e0297523 (2024).

35. Ostrem, J. M., Peters, U., Sos, M. L., Wells, J. A. & Shokat, K. M. K-Ras(G12C) inhibitors allosterically control GTP affinity and effector interactions. Nature 503, 548–551 (2013).

36. de Langen, A. J. et al. Sotorasib versus docetaxel for previously treated non-small-cell lung cancer with KRASG12C mutation: a randomised, open-label, phase 3 trial. Lancet 401, 733–746 (2023).

37. Holderfield, M. et al. Concurrent inhibition of oncogenic and wild-type RAS-GTP for cancer therapy. Nature 629, 919–926 (2024).

38. O’Reilly, E. M. et al. Daraxonrasib or chemotherapy in previously treated metastatic pancreatic cancer. N. Engl. J. Med. (2026).

39. Lu, A. et al. Engineered protein circuits for cancer therapy. bioRxiv 2025.04.16.647665 (2025).

40. Gray, D. C., Mahrus, S. & Wells, J. A. Activation of specific apoptotic caspases with an engineered small-molecule-activated protease. Cell 142, 637–646 (2010).

41. Orehek, S. et al. Cytokine-armed pyroptosis induces antitumor immunity against diverse types of tumors. Nat. Commun. 15, 10801 (2024).

42. Chambers, J. S., Brend, T. & Rabbitts, T. H. Cancer cell killing by target antigen engagement with engineered complementary intracellular antibody single domains fused to pro-caspase3. Sci. Rep. 9, 8553 (2019).

43. Vlahos, A. E. et al. Protease-controlled secretion and display of intercellular signals. Nat. Commun. 13, 912 (2022).

44. Xia, S. et al. Synthetic protein circuits for programmable control of mammalian cell death. Cell 187, 2785–2800.e16 (2024).

45. Chung, H. K. et al. A compact synthetic pathway rewires cancer signaling to therapeutic effector release. Science 364, eaat6982 (2019).

46. Zou, X., Zhao, C., Beier, K. T.Kang, C.-Y. & Lin, M. Z. Rewiring oncogenic signaling to RNA vector replication for the treatment of metastatic cancer (2025).

47. Senn, G. V., Nissen, L. & Benenson, Y. Synthetic gene circuits that selectively target RAS-driven cancers. Elife 14 (2026).

48. Watson, J. L. et al. De novo design of protein structure and function with RFdiffusion. Nature 620, 1089–1100 (2023).

49. Pacesa, M. et al. One-shot design of functional protein binders with BindCraft. Nature 646, 483–492 (2025).

50. Liu, B. et al. Design of high-specificity binders for peptide-MHC-I complexes. Science 389, 386–391 (2025).

51. Yang, J. et al. Active learning-assisted directed evolution. Nat. Commun. 16, 714 (2025).

52. Hoof, I. et al. NetMHCpan, a method for MHC class I binding prediction beyond humans. Immunogenetics 61, 1–13 (2009).

53. Nilsson, J. B. et al. Accurate prediction of HLA class II antigen presentation across all loci using tailored data acquisition and refined machine learning. Sci. Adv. 9, eadj6367 (2023).

54. Chen, B. et al. Predicting HLA class II antigen presentation through integrated deep learning. Nat. Biotechnol. 37, 1332–1343 (2019).

55. Wolfsberg, E. et al. Machine-guided dual-objective protein engineering for deimmunization and therapeutic functions. Cell Syst. 16, 101299 (2025).

56. Cheng, Q. et al. Selective organ targeting (SORT) nanoparticles for tissue-specific mRNA delivery and CRISPR-Cas gene editing. Nat. Nanotechnol. 15, 313–320 (2020).

57. Wang, X. et al. Preparation of selective organ-targeting (SORT) lipid nanoparticles (LNPs) using multiple technical methods for tissue-specific mRNA delivery. Nat. Protoc. 18, 265–291 (2023).

